# HORI-EN: Atomic-level energetic profiling and higher-order network identification in protein structures

**DOI:** 10.64898/2026.03.29.715065

**Authors:** Sarthak Joshi, Ramanathan Sowdhamini

**Affiliations:** National Centre for Biological Sciences (NCBS), Tata Institute of Fundamental Research (TIFR), Bangalore, Karnataka, India 560065; Molecular Biophysics Unit (MBU), Indian Institute of Science (IISc), Bangalore, Karnataka, India 560012; Institute of Bioinformatics and Applied Biotechnology (IBAB), Bangalore, Karnataka, India 560100

**Keywords:** Protein stability, Energetic Profiling, Residue Interaction Network, Mutational Hotspots, Higher-Order Interactions, Knowledge-Based Potentials

## Abstract

**Motivation:** Characterizing atomic-level stability and cooperative interaction networks is essential for understanding protein function and evolution. However, existing tools often lack the precision to integrate detailed physicochemical energies with higher-order graph-theoretic analyses.

**Results:** We present HORI-EN, an updated implementation to the HORI framework, featuring hybrid energetic scoring (Physicochemical + Knowledge-Based) and a Normalized Interaction Score (NIS) based on cumulative distribution functions. HORI-EN identifies higher-order cliques of interacting residues, revealing cooperative stabilization networks. Validation on the SKEMPI v2 dataset demonstrates that HORI-EN shows discriminative performance in identifying mutational hotspots, achieving an ROC-AUC of 0.780 on the full dataset and 0.844 on a clean benchmark. Enrichment analysis indicates a 3.1-fold increase in precision for the top 1% of predictions. Furthermore, analysis of the residue interaction network recovers 77.4% of non-contacting hotspots by identifying one-hop bridging interactions to the partner chain. Beyond hotspot prediction, HORI-EN distinguishes native structures from decoys and captures conserved energetic signatures in evolutionary case studies of serine proteases and lipases.

**Availability and Implementation:** The web server is freely available at https://caps.ncbs.res.in/HORI-EN and source code is available at https://github.com/thesixeyedknight/HoriPy.

## Introduction

The thermodynamic stability and functional specificity of biological macromolecules are governed by a delicate balance of non-covalent interactions. Since the foundational thermodynamic hypothesis of Anfinsen, it has been understood that the native three-dimensional structure of a protein is determined by the global minimization of free energy (Anfinsen 1973). This energy landscape is sculpted by opposing forces: the hydrophobic effect, which drives the formation of the compact core (Kauzmann 1959, Chothia 1974), and specific directional interactions such as hydrogen bonds (Pauling, Corey, and Branson 1951) and salt bridges, that provide geometric specificity and structural rigidity (Perutz 1978).

Quantifying these interactions remains a central challenge in structural biology. Early computational pioneers, such as Levitt and Lifson (Levitt and Lifson 1969), established that physics-based energy minimization could approximate protein conformations, while Miyazawa and Jernigan (Miyazawa and Jernigan 1985) demonstrated that statistical potentials derived from known structures could capture solvent-averaged contact energies. However, accurate energetic profiling is often hampered by the complexity of modeling environmental effects, particularly the local dielectric constant, which varies significantly between the protein core and the solvent-exposed surface (Warshel and Russell 1984, Burley and Petsko 1985). While recent methods have improved dielectric estimations (Pike and Nanda 2015), many standard tools still rely on bulk constants that oversimplify the electrostatic landscape.

Beyond pairwise energetics, there is increasing recognition that proteins function as cooperative networks rather than collections of isolated contacts. Graph-theoretic approaches have revealed that residues form “small-world” networks (Vendruscolo *et al*. 2002), where specific clusters or “cliques” of residues coordinate to stabilize active sites or transmit allosteric signals. Modern tools such as RING 2.0 (Piovesan, Minervini, and Tosatto 2016) and Arpeggio (Jubb *et al*. 2017) have made significant strides in visualizing these interaction networks. However, these tools often prioritize topological connectivity over detailed energetic scoring, or lack a unified framework that simultaneously accounts for atomic-level physics, statistical likelihood, and higher-order network structures.

To address this gap, HORI (Higher Order Residue Interactions) was previously developed, a web server and computational framework for analyzing protein interaction topologies (Sundaramurthy *et al*. 2010). While effective for identifying spatial clusters, the original implementation focused primarily on geometric contact mapping. In the intervening years, the explosion of high-resolution structural data and the availability of large-scale affinity benchmarks, such as SKEMPI v2 (Jankauskaitė *et al*. 2019), have necessitated a more robust, physically grounded approach.

In this study, we present HORI-EN (Hori-Energy Network), an extension of the original framework designed for structural and energetic analysis (Fig. 1A). HORI-EN integrates a hybrid scoring function that combines Knowledge-Based Potentials (KBP) with detailed physicochemical energies, utilizing the Pike-Nanda environment-dependent dielectric model (Pike and Nanda 2015) to model local electrostatic variations. Furthermore, we introduce the Normalized Interaction Score (NIS), a cumulative distribution function (CDF) based metric that enables the rigorous ranking of interaction quality across different physical types (Fig 1B). HORI-EN also identifies higher-order cliques of interacting residues, revealing cooperative stabilization networks (Fig. 1C2). We evaluate HORI-EN by validating its ability to predict mutational hotspots in the SKEMPI v2 dataset and distinguishing native structures from decoys with high discriminative performance. Finally, we apply the tool to evolutionary case studies of serine proteases and lipases, revealing conserved energetic signatures that persist despite extensive sequence divergence.

**Figure 1.**
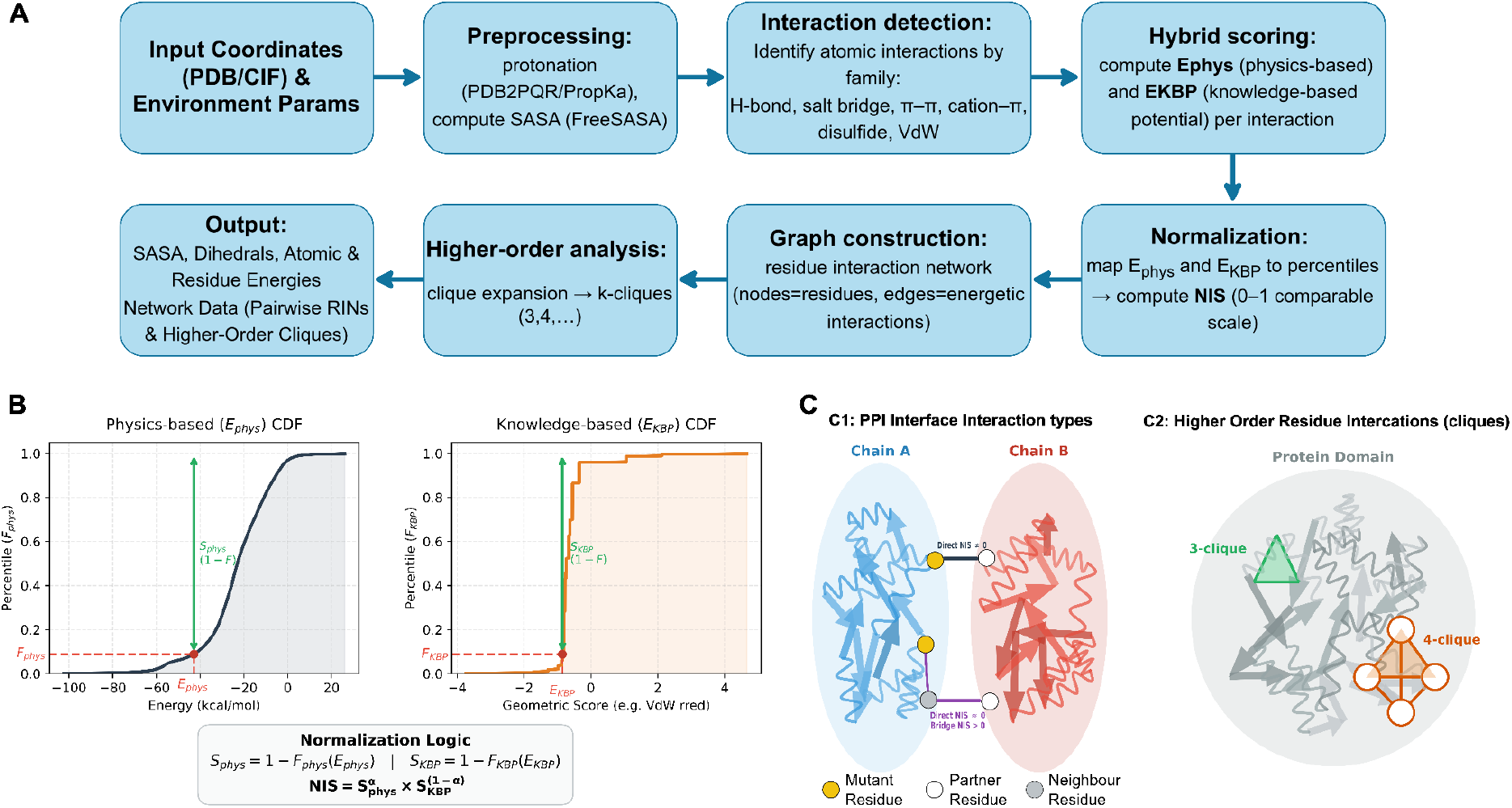
HORI-EN workflow, Normalized Interaction Score (NIS), and higher-order residue interaction networks. **(A)** End-to-end pipeline. Input protein coordinates (PDB/mmCIF) are preprocessed (protonation/hydrogens via PDB2PQR/PropKa; SASA via FreeSASA), followed by detection of atomic interaction types (H-bond, salt bridge, π–π, cation–π, disulfide, VdW). Each interaction is assigned a hybrid energetic description: a physicochemical term (E_phys) and a knowledge-based potential term (E_KBP). These are then normalized to a common 0–1 scale (NIS), used to build a residue interaction graph (nodes = residues; edges = energetically supported interactions), and expanded into higher-order residue cliques (k-cliques, k ≥ 3). The method outputs per-structure and per-residue energetic summaries and network/clique representations. **(B)** NIS computation. E_phys and E_KBP are mapped to empirical percentiles (CDFs) derived from background distributions, inverted to obtain S_phys = 1 − F_phys(E_phys) and S_KBP = 1 − F_KBP(E_KBP) (so larger values indicate more favorable interactions), and combined as a weighted geometric mean, NIS = S_phys^α × S_KBP^(1−α). **(C1)** Interface context for residue scoring: Direct contributions arise from contacts to the partner chain; bridge (one-hop) contributions arise when a residue connects to the partner chain through a single intermediate neighbor in the residue interaction network. **(C2)** Higher-order residue interactions: illustrative 3-clique and 4-clique motifs extracted from the residue interaction graph, representing cooperative interaction modules beyond pairwise contacts.

## System and Methods

### Training Dataset and Knowledge-Based Potential Generation

To derive robust statistical potentials, we utilized a non-redundant subset of the Protein Data Bank (PDB) generated via the PISCES culling server (Wang and Dunbrack 2003). The dataset was filtered to include only X-ray structures with a resolution better than 2.0 Å, an R-factor ≤ 0.25, and a maximum pairwise sequence identity of 30% to strictly eliminate homologous redundancy. Chains with chain breaks or lengths outside the 40–10,000 residue range were excluded. This yielded a training corpus of 9,528 unique protein chains (dataset pc30_res2.0_d2025_05_12).

We extracted geometric distributions for six interaction families: Hydrogen Bonds (distance, angle), π-π stacking (centroid distance, plane angle), Cation-π (cation-centroid distance), Salt Bridges (distance), Disulfide bonds (χ_*SS*_, χ_1_ dihedrals), and Van der Waals contacts (reduced radius). Reference states were computed using a distance-dependent random mixing model to normalize observed frequencies against background probabilities.

### Validation Datasets

To assess the discriminative power and predictive accuracy of HORI-EN, we employed two independent datasets, SKEMPI v2 database and Titan High-Resolution decoy set.

#### Affinity and Hotspot Classification (SKEMPI v2)

We utilized the SKEMPI v2 database (Jankauskaitė *et al*. 2019) to evaluate association between energetic scores and experimental binding free-energy changes (ΔΔ*G*). The dataset was filtered to retain single-point mutations with available dissociation constants (*K*_*d*_). We used the officially provided “cleaned” PDB structures and mapping files to ensure accurate alignment.

The dataset was stratified into a “Full Dataset” and a “Clean Benchmark” for rigorous testing. In the Clean Benchmark, mutational hotspots were defined as residues where ΔΔ*G* ≥ 2. 0 kcal/mol, and neutral sites as |ΔΔ*G*| ≤ 0. 5 kcal/mol; mutations with intermediate values (0. 5 < |ΔΔ*G*| < 2. 0) were excluded.

For each mutation site, we calculated a cumulative interaction score representing the sum of NIS contributions from all interface partners. This cumulative score aggregates two distinct components: (i) Direct NIS, derived from residues in direct atomic contact, and (ii) Indirect (Bridge) NIS, derived from one-hop interactions where the mutated residue is connected to the partner chain via a single intermediate residue in the interaction network (Fig. 1C1).

#### Structural Decoy Discrimination (Titan HR)

To evaluate whether HORI-EN distinguishes experimentally determined native structures from near-native misfolded models, we analyzed the Titan High-Resolution (Titan HR) decoy set (Rajgaria, McAllister, and Floudas 2006). For each target, HORI-EN computed structure-level summary metrics derived from the energetic and interaction-profile outputs, including (i) the mean physicochemical energy per residue (total *E*_*phys*_ divided by sequence length), (ii) the number (or rate) of high-energy knowledge-based potential (KBP) violations, defined as interactions with *E*_*KBP*_ > 3. 0 *kT*, and (iii) a hydrophobic exposure ratio, defined as the fraction of solvent-accessible surface area attributable to hydrophobic residues (computed from FreeSASA using the same preprocessing pipeline).

Models with severe steric clashes were excluded using a mean physicochemical energy threshold (> 50.0 kcal/mol/residue) to focus the evaluation on decoys that are not trivially separable by clashes. Discriminative performance was quantified using ROC-AUC by treating native structures as positives and decoys as negatives for each metric.

#### Evolutionary Case Studies

Two specific systems were selected to analyze energetic conservation amidst sequence divergence: (1) The Serine Protease family, comparing Trypsin (PDB: 2PTN), Chymotrypsin (PDB: 4CHA), and Elastase (PDB: 3EST); and (2) The α/β hydrolase superfamily, comparing a canonical lipase (PDB: 1TGL) with a rewired evolutionary intermediate (PDB: 1ETH).

## Algorithm

### Preprocessing

All input structures (PDB/mmCIF) are pre-processed to add hydrogen atoms and determine protonation states at pH 7.0 using PropKa (Olsson *et al*. 2011) via the PDB2PQR framework (Jurrus *et al*. 2018). The framework simultaneously repairs the structure by reconstructing missing heavy atoms and optimizing the hydrogen bond network to resolve steric conflicts. Solvent-Accessible Surface Area (SASA) is computed using the FreeSASA library (Mitternacht 2016) with a standard probe radius of 1.4 Å.

### Geometric Parsing

HORI-EN parses 3D atomic coordinates to infer molecular topology. Unlike tools that rely solely on Euclidean distance cutoffs (e.g., RING 2.0), HORI-EN employs a hybrid geometric-chemical definition logic. Atomic types are mapped to the AMBER force field nomenclature to assign radii, charges, and hybridization states.

### Hybrid Energetic Scoring

HORI-EN integrates physicochemical mechanics with statistical likelihoods into a unified framework.

#### Physicochemical Energy (*E*_*phys*_)

Electrostatic interactions (Salt Bridges, Hydrogen Bonds) are calculated using Coulomb’s law. To address the limitations of bulk dielectric constants in heterogeneous protein environments, we implemented the Pike-Nanda model (Pike and Nanda 2015). This method computes a local environment-dependent dielectric constant (*ϵ*_*loc*_) based on the local polarizability density (ρ_α_) within a 9.0 Å sphere of the interacting atoms:

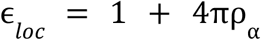

Van der Waals forces are calculated using a 6-12 Lennard-Jones potential. Specialized formulations are applied for π-π interactions (damped by angular deviation from parallel or T-shaped optimality) and Cation-π interactions.

#### Knowledge Based Potentials (*E*_*KBP*_)

For every interaction type *i* and geometric feature *x* (e.g., angle *θ*), we compute a pseudo-energy potential of mean force (PMF) using the inverse Boltzmann principle:

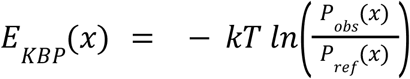

where *P*_*obs*_ is the observed frequency in the training set and *P*_*ref*_ is the reference state frequency.

### Normalised Interaction Score (NIS)

Raw energy values vary wildly across interaction types (e.g., a salt bridge is orders of magnitude stronger than a VdW contact), making cross-comparison difficult. We introduce the Normalized Interaction Score (NIS), a probabilistic metric based on Cumulative Distribution Functions (CDF) (Fig. 1B).

For a given interaction with physical energy *E*_*phys*_ and KBP energy *E*_*KBP*_, we map values to their empirical percentiles (*F*) derived from the background distribution of the training corpus. The raw scores are inverted such that 1.0 represents the most favorable interaction:

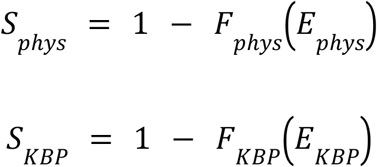

The final NIS is computed as the geometric mean weighted by a factor of *α* (typically 0.5):

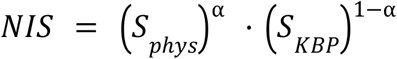

This formulation ensures that a high NIS score (approaching 1.0) strictly indicates an interaction that is both physically stable and statistically enriched in native structures.

### Higher-Order Residue Interaction (HORI) Networks

Beyond pairwise contacts, HORI-EN identifies cooperative stability networks. The algorithm constructs an adjacency graph where residues are nodes and valid energetic interactions are edges. We employ a recursive clique-finding algorithm that iteratively expands *k*-cliques to (*k* + 1)-cliques until convergence, effectively isolating clustered hubs of 3-body, 4-body, and higher-order residue interactions essential for structural cores and active sites.

## Implementation

### Core Framework and Architecture

HORI-EN is implemented as a modular Python-3 library designed for extensibility and high-throughput analysis. The software architecture is organized into four distinct functional layers: structure parsing, geometric topology, energetic scoring, and graph-theoretic analysis. The parsing layer handles standard PDB and mmCIF inputs, integrating PropKa via the PDB2PQR framework to determine protonation states at a user-specified pH. The topology layer utilizes NumPy for vectorized distance calculations, constructing neighbor lists and spatial grids to efficiently identify atomic contacts.

The energetic scoring layer is further subdivided into physicochemical and statistical modules. The physicochemical module implements the Pike-Nanda dielectric model and AMBER-based force field calculations, while the statistical module manages Knowledge-Based Potentials (KBP) via efficient lookups in pre-computed interaction matrices. A dedicated scoring engine, the NIS module, normalizes these diverse energy terms using cumulative distribution functions. Finally, the graph-theoretic layer employs the multiprocessing library to parallelize the recursive expansion of pairwise contacts into higher-order cliques. Computationally intensive tasks, such as solvent-accessible surface area (SASA) determination, are offloaded to the FreeSASA C-library bindings for performance.

### Web Server and User Interface

We developed a web interface using the Flask microframework in order to make the tool readily accessible to the broader community. The server employs a non-blocking asynchronous design, where user submissions are routed to an in-memory First-In-First-Out (FIFO) job queue and processed by background daemon threads. This architecture prevents request timeouts during computationally demanding analyses of large macromolecular assemblies.

Data persistence is managed via MongoDB, which serves strictly as a temporary cache for job metadata and analysis results. Processed data is retained for a period of one month to facilitate retrieval and sharing via unique UUIDs, after which it is automatically purged to ensure storage efficiency. The client-side interface is built with Bootstrap 5 for responsive design and integrates specialized JavaScript libraries for interactive visualization: 3Dmol.js for hardware-accelerated molecular rendering, Cytoscape.js for the dynamic exploration of higher-order interaction networks, and Chart.js for statistical profiling of energetic distributions.

### Availability

The HORI-EN source code is freely available to the academic community under the GNU GPLv3 license and is hosted on GitHub (https://github.com/thesixeyedknight/HoriPy). The web server can be accessed freely without login requirements at https://caps.ncbs.res.in/HORI-EN. Documentation is provided in the repository to facilitate local installation and reproduction of results.

## Discussion

### The Normalized Interaction Score (NIS) as a Hotspot Classifier

Raw energy values often vary by orders of magnitude across interaction types (e.g., a salt bridge vs. a Van der Waals contact), rendering it difficult to rank the relative importance of different residues. A feature of HORI-EN is the Normalized Interaction Score (NIS), which maps these heterogeneous values onto a unified probabilistic scale (0 to 1) using Cumulative Distribution Functions.

In validation against the SKEMPI v2 dataset (Jankauskaitė *et al*. 2019), NIS was assessed for the identification of mutational hotspots (ΔΔ*G* ≥ 2. 0 kcal/mol). On the Full Dataset (please see methods for details), the method yielded an ROC-AUC of 0.78 (Fig. 2B), with strong early enrichment among top-ranked predictions (Fig. 2C). When evaluated against the Clean Benchmark (excluding ambiguous ΔΔ*G* values), the discriminative performance improved to an ROC-AUC of 0.844. Restricting the analysis to the Alanine Scanning subset resulted in an ROC-AUC of 0.859. The aggregate score of all atomic interactions involving a specific residue outperformed the maximum single-interaction score, suggesting that a residue’s contribution to stability is cumulative (Fig. 2A).

**Figure 2.**
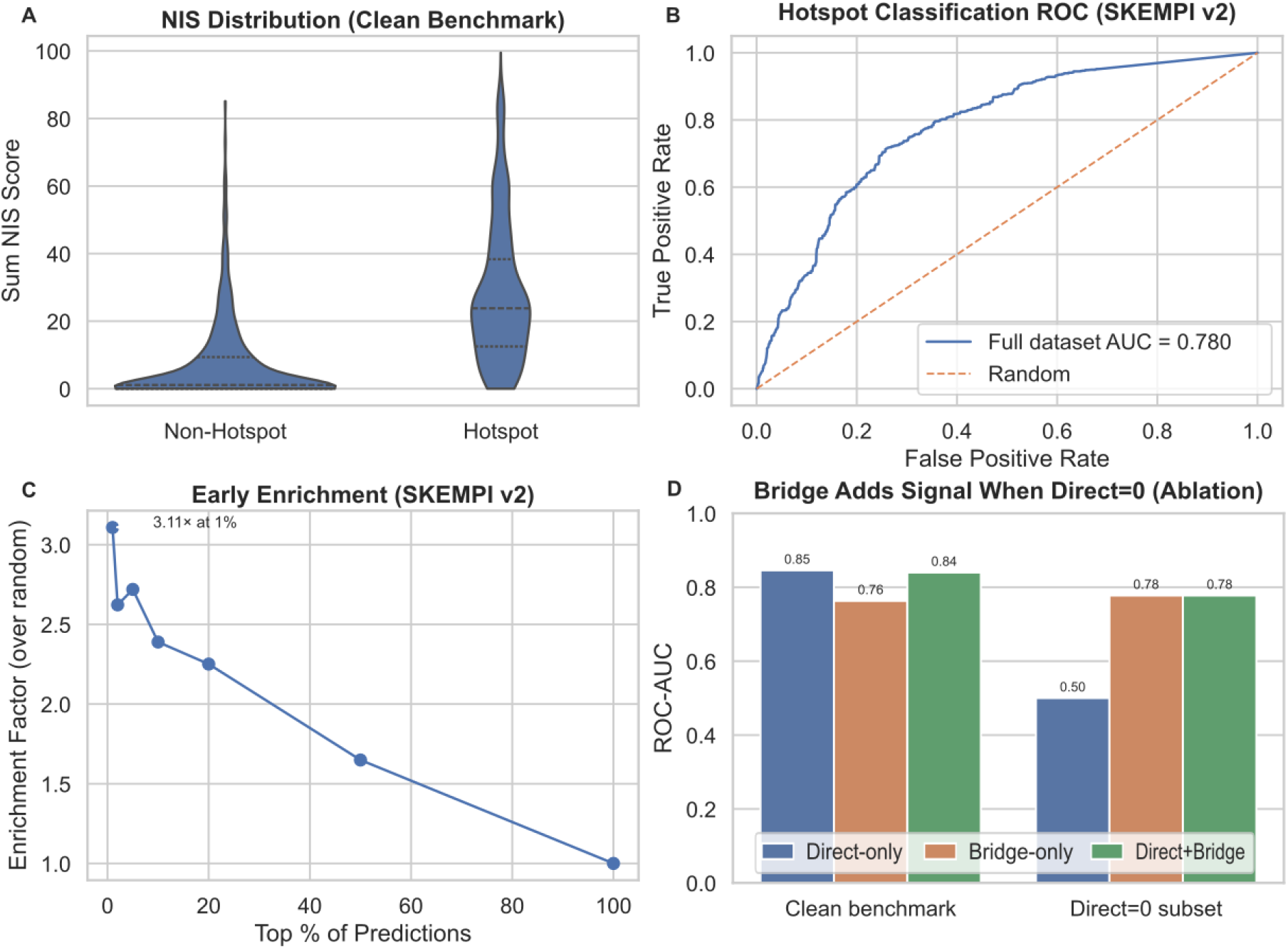
HORI-EN hotspot prediction on SKEMPI v2 and contribution of network-mediated (bridge) interactions. **(A)** Violin plots of residue-level cumulative interaction strength at mutation sites, quantified as the Sum NIS score (sum of all partner-facing NIS contributions at the site). The plot uses the clean benchmark definition (hotspots: ΔΔG ≥ 2.0 kcal/mol; non-hotspots: |ΔΔG| ≤ 0.5 kcal/mol; intermediate mutations excluded) and shows that hotspots are shifted toward substantially higher Sum NIS. **(B)** ROC curve for hotspot classification on the full SKEMPI v2 dataset using Sum NIS as the predictor (AUC = 0.780; diagonal line indicates random performance). **(C)** Early-enrichment analysis for the same ranking: hotspots are enriched among the top-scoring residues, with a 3.11× enrichment in the top 1% of predictions relative to the dataset baseline. **(D)** Ablation demonstrating that one-hop bridge signal adds predictive power specifically when direct interface contacts are absent. Bars show ROC-AUC for Direct-only, Bridge-only, and Direct+Bridge scoring, evaluated on the clean benchmark (Direct-only 0.85; Bridge-only 0.76; Direct+Bridge 0.84) and on the subset of mutation sites with Direct = 0 (Direct-only 0.50; Bridge-only 0.78; Direct+Bridge 0.78). Bridge score is computed as the sum of indirect interaction terms (indirect_inner_nis + indirect_outer_nis), and the combined model uses Direct NIS + Bridge NIS.

A subset of hotspots in the SKEMPI v2 dataset exhibits no direct interchain interactions (Direct NIS = 0). Under the structural region scheme of (Levy 2010), these sites are enriched in Support (39%) and Interior (25%) regions rather than the Core interface. This is consistent with the finding of secondary shell hotspots reported by (Parvathy *et al*. 2025). Solvent accessibility is consistent with this classification: these residues show minimal change upon complex formation (mean Δ*SASA* ≈ 15 Å^2^) compared to direct-contact hotspots (mean Δ*SASA* ≈ 86 Å^2^). Despite the lack of direct physical contact, these residues contribute significantly to binding affinity (Fig. 1C1). We applied a network analysis to these sites using the residue interaction network generated by HORI-EN. By examining the immediate neighborhood of these residues, we identified bridging interactions. These are cases where the mutated residue connects to the partner chain through a single intermediate residue. This approach recovered 77.4% of the non-contacting hotspots (Fig. 2D). These results indicate that a substantial fraction of experimentally defined hotspots can be connected to the partner chain through short interaction paths even in the absence of direct atomic contacts, thereby demonstrating the potential to rescue performance on these challenging targets from random to discriminative (AUC 0.78) levels.

### Decoy Discrimination and the Energetic Landscape

A fundamental challenge in structural bioinformatics is distinguishing biologically relevant interactions from incidental geometric contacts. While distance-based contact maps are computationally inexpensive, they treat all neighbors within a cutoff as equal. Consequently, they often fail to capture the thermodynamic subtleties (or “frustrations”) that destabilize misfolded structures (Frauenfelder, Sligar, and Wolynes 1991). Our validation on the Titan HR decoy set demonstrates that HORI-EN addresses this by enforcing strict physicochemical constraints alongside geometric ones.

Our analysis reinforces the “Hydrophobic Collapse” hypothesis—the concept that the burying of non-polar surface area is the dominant driving force in the initial stages of protein folding (Kauzmann 1959, Dill 1990). This is statistically evidenced by the near-perfect discrimination between native structures and decoys using the Hydrophobic Exposure Ratio (AUC = 1.00). Native structures consistently exhibit a tighter packing of hydrophobic residues compared to decoys, a feature that geometric contact maps often overlook (Novotný, Rashin, and Bruccoleri 1988, Nabuurs *et al*. 2006).

However, the energy landscape of a protein is more nuanced than simple compaction and exhibits a “funnel-shaped” landscape: while native structures occupy a deep energetic minimum distinct from decoys (Fig. 3A), the metrics showed only a weak linear correlation with RMSD (ρ ∼ 0. 1 − 0. 3) (Fig. 3D). This observation aligns with the Principle of Minimal Frustration (Bryngelson and Wolynes 1987), which posits that native proteins are evolved to minimize conflicting interactions. This is quantitatively supported by our finding that High-Energy KBP Violations (> 3.0 kT) are powerful discriminators (AUC = 0.95) (Fig. 3B), whereas the permissive KBP contact-count metric (U > −1.5) shows the opposite trend (Fig. 3C). Thus, HORI-EN identifies native-like quality not merely by the maximization of favorable contacts, but by the conspicuous absence of severe energetic conflicts. This is a distinction critical for accurate structural assessment.

**Figure 3.**
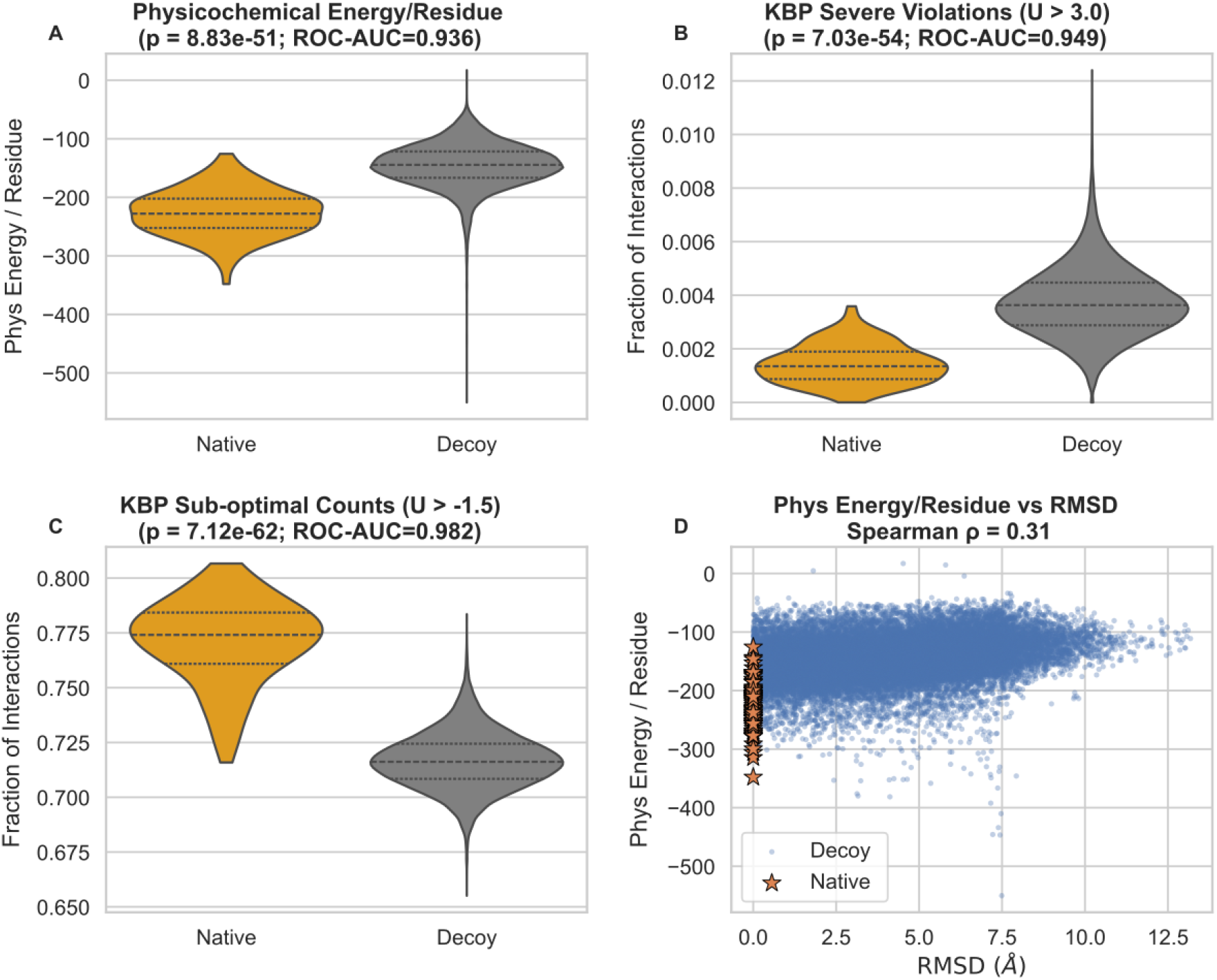
Native–decoy discrimination in Titan HR highlights the role of energetic conflicts versus “favorable-contact” abundance. **(A)** Distribution of mean physicochemical energy per residue for native structures and decoys after filtering out trivially clashing models (Methods). Native structures occupy a more favorable energy range (Mann–Whitney U p = 8.83×10^−51^; ROC-AUC = 0.936, with “more native-like” defined by lower energy). **(B)** Distribution of the fraction of severe KBP violations (U > 3.0 kT). Natives exhibit fewer high-energy conflicts than decoys (p = 7.03×10^−54^; ROC-AUC = 0.949). **(C)** Distribution of the fraction of sub-optimal / non-strongly-favorable KBP interactions (U > −1.5 kT). This metric shows the opposite trend to Panel B (p = 7.12×10^−62^; ROC-AUC = 0.982), consistent with decoys exhibiting a relative excess of very favorable interactions (U ≤ −1.5) despite being non-native, illustrating that *maximizing “good contacts” alone is not sufficient* to identify native-like structures, whereas the depletion of severe conflicts (Panel B) provides a mechanistically interpretable quality signal. **(D)** Scatter of physicochemical energy per residue versus RMSD for decoys, with natives highlighted (Spearman ρ = 0.31). The weak correlation indicates that energetic quality and geometric similarity are only partially coupled, consistent with a funnel-like landscape where native structures occupy a distinct energetic basin.

### Energetic Conservation in Protein Evolution

Evolutionary theory suggests that while protein sequences diverge, the energetic features critical for function must be conserved. By applying HORI-EN to evolutionary case studies, we observed that energetic signatures often persist longer than sequence identity.

In the Serine Protease family, we compared Trypsin, Chymotrypsin, and Elastase (Fig 4A,B). Despite sequence identities dropping below 40%, the catalytic triad maintained a consistent total interaction energy of approximately -500 kcal/mol across all homologs (Fig 4B). This quantitative stability suggests that evolutionary pressure acts to preserve the thermodynamic profile of the active site scaffold, even as the surrounding sequence space drifts.

**Figure 4.**
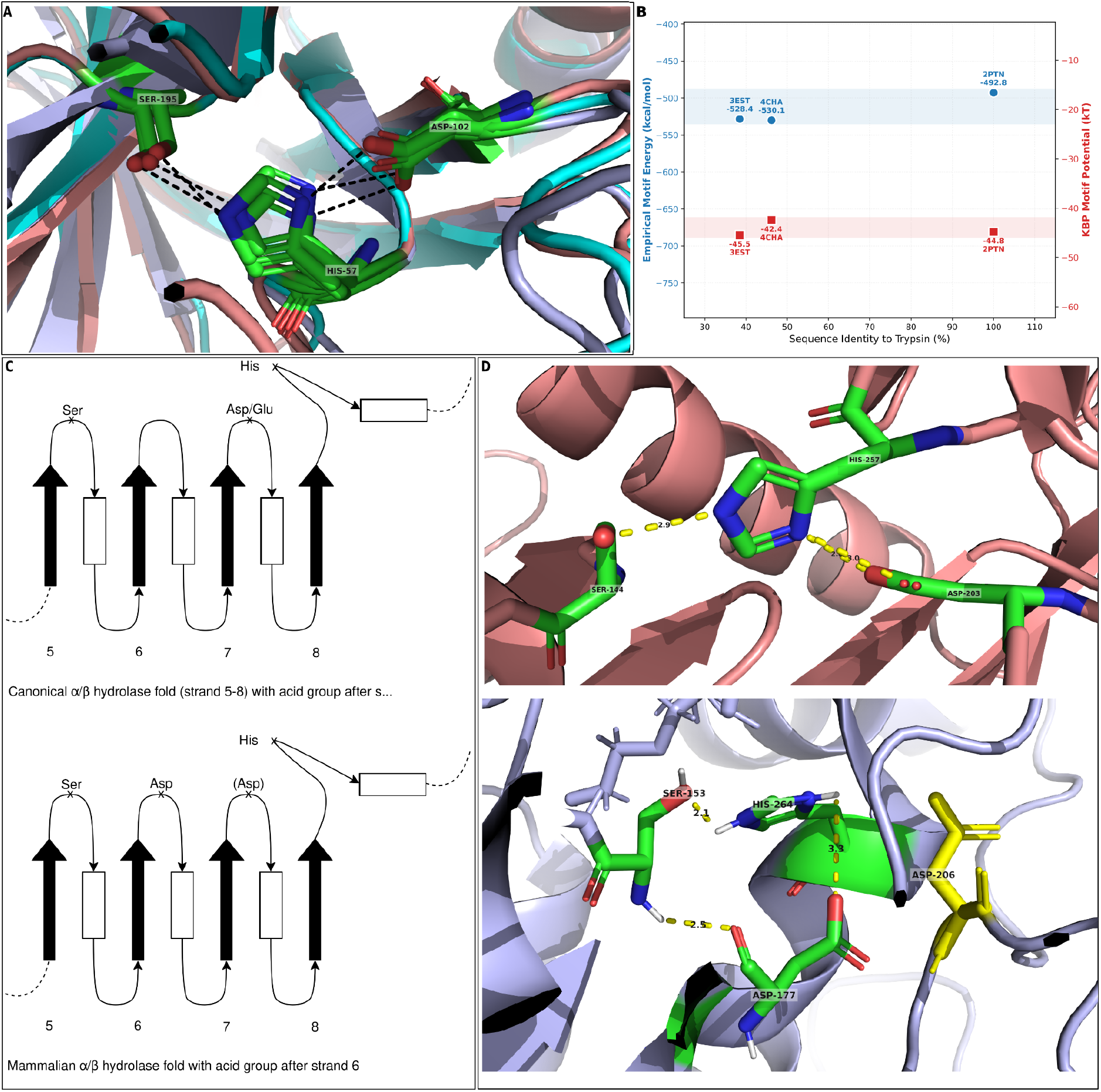
Conserved energetic signatures and catalytic-acid rewiring detected by HORI-EN. **(A)** Structural superposition of serine proteases Trypsin (2PTN), Chymotrypsin (4CHA), and Elastase (3EST) highlighting the catalytic triad region (His57, Asp102, Ser195; residue numbering shown on the representative model). The triad geometry and local interaction motif are preserved despite substantial sequence divergence. **(B)** Motif energetics plotted against sequence identity to trypsin for the serine protease triad module. Blue circles report the empirical/physicochemical motif energy (kcal/mol; left y-axis) and red squares report the knowledge-based (KBP) motif potential (kT; right y-axis) for 2PTN, 4CHA, 3EST. Shaded bands indicate the narrow range of motif energies across homologs, emphasizing energetic conservation despite reduced pairwise sequence identity. **(C)** Schematic of catalytic-acid migration in the α/β-hydrolase superfamily, contrasting the canonical placement of the acidic residue after β-strand 7 with a rewired placement after β-strand 6. **(D)** Active-site closeups illustrating functional rewiring in α/β hydrolases comparing 1TGL (top; canonical lipase) and 1ETH (bottom; rewired intermediate). In 1TGL the canonical catalytic triad is shown (Ser144–His257–Asp203). In 1ETH the Ser–His pair is conserved (Ser153–His264) but the functional acidic residue is located at the rewired position (Asp177), while the residue at the canonical position (Asp206) is retained as a non-functional structural relic. Distances indicate key hydrogen-bond contacts supporting the catalytic configuration.

Furthermore, the tool successfully tracked active site migration in the α/β Hydrolase superfamily (Schrag, Winkler, and Cygler 1992). By comparing a canonical lipase (1TGL) with an evolutionary intermediate (1ETH), HORI-EN quantified the shift of the catalytic acid from strand 7 to strand 6 (Fig. 4C,D). The energetic scoring correctly identified the residue at the original position as a “structural relic” (possessing low interaction energy) and the residue at the new position as the functional anchor (possessing high interaction energy). This capability to detect “energetic rewiring” highlights the potential of HORI-EN for annotating function in proteins where sequence homology is ambiguous.

## Limitations and Future Directions

The current implementation of HORI-EN relies on rigid-body structural approximations. The Pike-Nanda dielectric model and KBP scoring are calculated on static coordinates, which effectively treats the protein as a frozen snapshot. This approach may overlook entropic contributions arising from side-chain flexibility and backbone fluctuations.

Future iterations of the framework will aim to incorporate ensemble-based scoring. Rather than scoring a single static PDB file, this approach involves generating a diverse set of conformations (an ensemble) via Molecular Dynamics simulations or NMR data. By averaging energetic scores across this ensemble, the tool could better account for the conformational dynamics and entropic factors that are intrinsic to protein stability (Karplus and McCammon 2002). Additionally, while the integration of PropKa mitigates errors in protonation states (Olsson *et al*. 2011), the accuracy of electrostatic terms remains sensitive to the resolution of the input structure.

